# Spatial odor discrimination in hawkmoth, *Manduca sexta* (L.)

**DOI:** 10.1101/2020.07.08.194126

**Authors:** P. Kalyanasundaram, M. A. Willis

**Author notes:** Contact: Parthasarathy Kalyanasundaram, Department of Biology, Case Western Reserve University, Cleveland, OH 44107-7080.

## Abstract

Flying insects track turbulent odor plumes to find mates, food and egg-laying sites. To maintain contact with the plume, insects are thought to adapt their flight control according to the distribution of odor in the plume using the timing of odor onsets and intervals between odor encounters. Although timing cues are important, few studies have addressed whether insects are capable of deriving spatial information about odor distribution from bilateral comparisons between their antennae in flight. The proboscis extension reflex (PER) associative learning protocol, originally developed to study odor learning in honeybees, was modified to show hawkmoths, *Manduca sexta*, can discriminate between odor stimuli arriving on either antenna. We show moths discriminated the odor arrival side with an accuracy of >70%. The information about spatial distribution of odor stimuli is thus available to moths searching for odor sources, opening the possibility that they use both spatial and temporal odor information.

## 1. Introduction

Animals with bilaterally symmetrical sensory structures (i.e., eyes, ears, nostrils, antennae, etc.) assess information by either combining and comparing bilateral inputs from sequential time points (temporal sampling), or comparing bilateral inputs at the same time point (spatial sampling) (Dusenbery, 1992; Fraenkel and Gunn, 1961; Schone and Strausfeld, 1984). Several studies have shown that a variety of animals use bilaterally sampled inputs from their olfactory organs to orient towards odor sources (Borst and Heisenberg, 1982; Catania, 2013; Duistermars et al., 2009; Gardiner and Atema, 2010; Khan et al., 2012; Kraus-Epley and Moore, 2002; Louis et al., 2008; MARTIN, 1965; Page et al., 2011; Parthasarathy and Bhalla, 2013; Porter et al., 2007; Rajan et al., 2006; Takasaki et al., 2012; Vonbekesy, 1964; Wasserman et al., 2012; Weissburg and Dusenbery, 2002). These studies employ a variety of techniques including genetic manipulations in controlled laboratory environments (Gomez-Marin et al., 2011; Louis et al., 2008), unilateral odor stimulation in tethered walking and flying preparations (Borst and Heisenberg, 1982; Duistermars et al., 2009; Takasaki et al., 2012), telemetric unilateral stimulation of freely swimming sharks(Gardiner and Atema, 2010), and experimentally manipulated individuals in naturalistic environments(Catania, 2013; Gardiner and Atema, 2010; Khan et al., 2012; Kraus-Epley and Moore, 2002; Parthasarathy and Bhalla, 2013; Rajan et al., 2006; Takasaki et al., 2012). Despite the separation between sensory organs ranging from less than a millimeter (Louis et al., 2008) to multiple centimeters (Gardiner and Atema, 2010), asymmetric stimulation usually caused asymmetric steering or the inability to maintain orientation to stimulus(Gardiner and Atema, 2010; Louis et al., 2008). Odor asymmetry was detected when presented as difference in odor concentration or onset timing (Gardiner and Atema, 2010; Parthasarathy and Bhalla, 2013; Rajan et al., 2006; Vonbekesy, 1964).

Previous studies assumed that bilaterally located sensors enhance spatial sampling. However, studies on plume tracking during flight typically assume that insects do not spatially sample odors, because turbulent odor plumes may not provide predictable directional cues leading to the source. Also, freely-flying plume trackers may move too fast for their nervous systems to process bilateral asymmetries and accordingly maneuver. Directional cues are thought to be provided by the wind or water flow detected visually (Emanuel and Dodson, 1979; Fry et al., 2009; Kennedy, 1940; Kennedy and Marsh, 1974) or via mechanosensation (Baker and Montgomery, 1999; Kulpa et al., 2015), and turning maneuvers underlying zigzagging trajectories are thought to be pre-programmed in the central nervous system and expressed in response to an attractive odor (Arbas et al., 1993; Kennedy, 1983). Bilateral sampling has been deemed unnecessary(Arbas et al., 1993).

Few studies have directly addressed if bilateral olfactory comparisons contribute to plume tracking in moths (Takasaki et al., 2012; Vickers and Baker, 1991). Experiments in which male *Heliothis virescens* (Fabricius) moths tracked a pheromone plume with one antenna removed showed minor changes in their flight trajectories and interestingly, removal of an antenna diminished successful plume tracking by roughly ∼50% relative to the intact controls (Vickers and Baker, 1991). This result was attributed to the loss of odor inputs and timing of odor onset resulting from moths with asymmetric antennae encountering the edges vs. the centerline of a laboratory-generated odor plume. The moth’s inability to make bilateral comparisons across two antennae was discounted (Vickers and Baker, 1991). In contrast, Takasaki et al. showed that walking silk moths, *Bombyx mori* (L.) alter their steering in response to asymmetric odor stimulation(Takasaki et al., 2012). However, they cautioned that walking and flying are very different modes of locomotion. Because flying insects move much faster than walking insects, they may use bilateral odor information differently (Takasaki et al., 2012). Indeed, the use of spatial sampling during plume tracking in flight remains an open question, even though odor plume tracking is a well-studied topic in flying moths and flies tracking attractive odors (Willis and Arbas, 1991; Cardé and Willis, 2008; Duistermars et al., 2009; Saxena et al., 2017).

Before we try to understand the role of spatial sampling in plume tracking during flight, we must first establish that moths are capable of discriminating between olfactory stimuli detected at one antenna *vs*. the other. We studied this question in the hawkmoth *M. sexta* using an associative learning paradigm based on the proboscis extension reflex (PER) conditioning developed for honey bees (Kuwabara, 1957). Building on previous studies that used PER to demonstrate associative learning in hawkmoths (Daly and Smith, 2000; Daly et al., 2001; Daly et al., 2008), we developed a behavior paradigm to probe the spatial odor representation in moths. In our study, trained moths discriminated the odor arrival side with an accuracy of >70%. Thus, moths can distinguished the odor arrival side and may be capable of extracting spatial information about the odor distribution while tracking plumes in flight.

## 2. Materials and methods

### (a) Animals

We used 4-day old female *M. sexta* (L.) moths reared in our laboratory and maintained on a 14:10 Light: Dark cycle. Experiments were conducted during the dark phase of the L:D cycle when plume tracking typically occurs in nature (Sasaki and Riddiford, 1984).

### (b) Experimental setup

#### Odor delivery system

Each antenna (left or right) received odor from a custom-built odor delivery system consisting of blank, empty glass vials and a glass vial containing an odor-soaked Whatman #1 filter paper strip (0.5 × 10 mm). The stimulus was produced by letting clean air (1.5 liters/min) flow through the glass vial containing saturated odor vapor, and odorized air was injected into the tube delivering odor to an antenna (Fig. 1a). Air was filtered using a charcoal filter and bubbled through water to humidify it before reaching the odor vials. Odor vials were changed every day and fresh odor loaded into the vial before every training session. During inter-trial intervals, air flowed through the blank vial to ensure constant flow of air onto the moth. Each antenna was inserted into separate independent odor delivery tubes (1.6 mm id, 3.2 mm od; Fisher scientific, USA.) to eliminate cross-contamination between antennae (Fig. 1a). An exhaust fan setup placed directly above the preparation, removed stimuli away from the room. Airflow in the olfactometer was gated by solenoid valves (Clippard Instrument Laboratory, Fairfield, OH, USA. ET-2M-12H) interfaced with a computer using a 2-channel relay controller (National Control Devices, Osceola, Missouri, USA. R210RS), driven by a custom MATLAB code. During a trial, odorized air was delivered to one antenna while the other received flow-speed-matched clean air from an empty blank vial.

**Figure 1.**
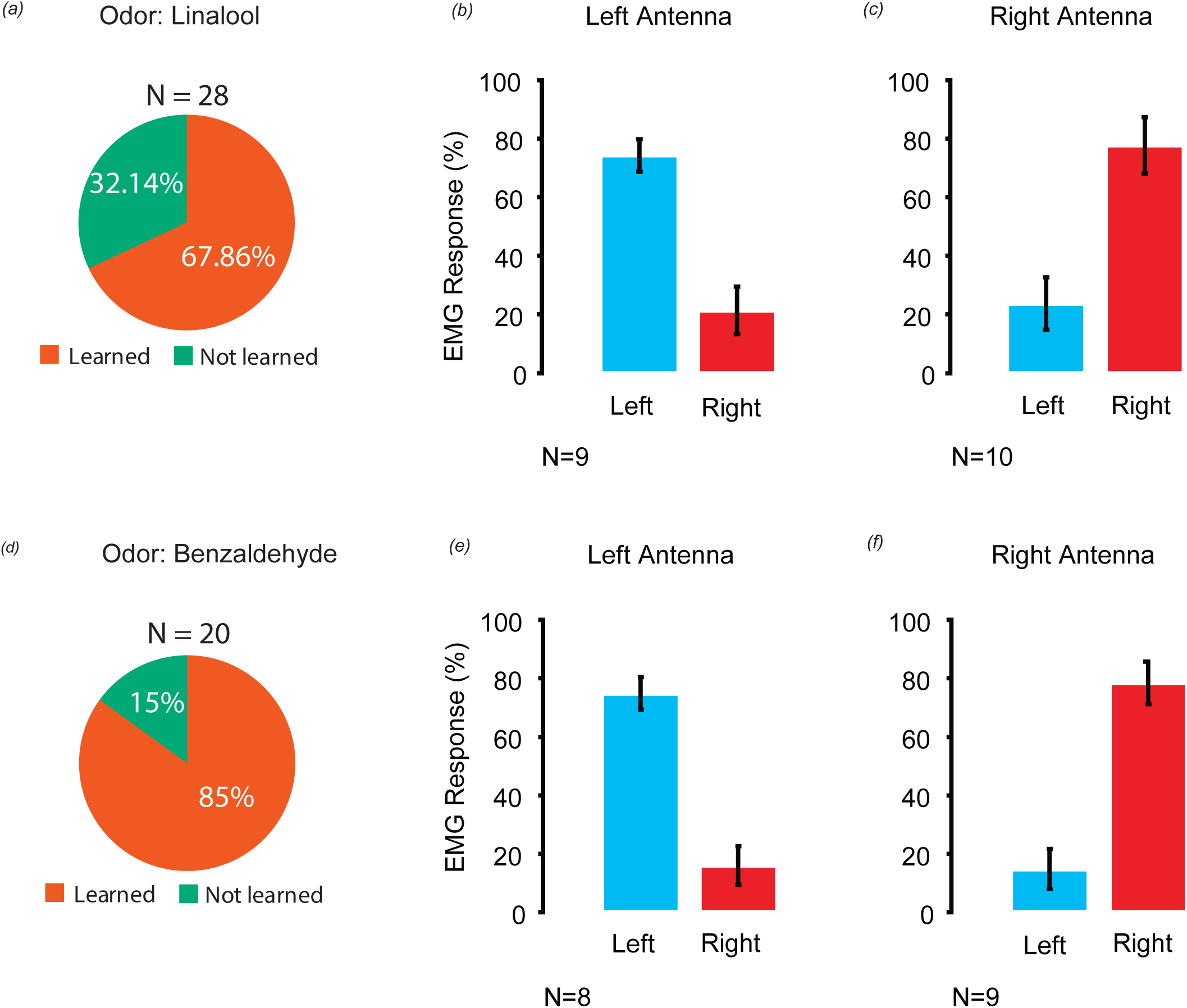
Behavioral setup and training protocol. (a) Schematic diagram of behavioral setup showing restrained moth with its straightened proboscis and odor delivery onto its antennae. Reward delivery port was placed at the distal tip of the proboscis. The feeding response was monitored using EMG activity of the suction pump muscles in the head. (b) Schematic diagram shows events during training and test protocols. Odor was presented for 5 s and 3 s after odor onset the reward valve was activated for 0.1 s. Moths were not rewarded during test session.

We used the odorants linalool and benzaldehyde (purity ≥ 99%, Sigma-Aldrich, USA) to serve as conditioned stimuli. Based on available literature on odor coding in antennal lobes, these two chemicals activate different elements of odor detection and processing circuitry (Bisch-Knaden et al., 2018; Martin et al., 2011; Saveer et al., 2012). We assumed that any observed associations would be general responses. These two compounds have been identified in scents released by night-blooming flowers on which *M. sexta* feed (Raguso et al., 2003a; Raguso et al., 2003b). They also attract and stimulate feeding in *M. sexta* (Reisenman et al., 2010; Riffell et al., 2009).

### (c) Training protocol

An intact moth was held immobile in a 6.4 cm long piece of 10 cm diam. copper tubing, with the head attached to the tube surface using soft dental wax (Fig. 1a). Reward (10% sucrose in water) was delivered to the tip of experimentally extended 7.5 cm long proboscis (i.e., proboscis was threaded through a 4 cm long, 0.4 cm diam. plastic tube). The tube was cut lengthwise to allow rapid placement of the proboscis. The distal tip of the proboscis was free to coil around the reward delivery tube. A pinch valve (161P010; West Caldwell, NJ, USA) was used to control the reward delivery duration. We monitored feeding attempts by recording electromyograms (EMGs) through a pair of Teflon-coated silver wires (bare dia., 0.127 mm; A-M systems, Sequim, WA, USA) (Daly and Smith, 2000) implanted in the cibarial pump muscle, which when activated draws nectar along the proboscis. Use of cibarial muscle EMGs to monitor associations in this preparation was developed because tethered *M. sexta* moths do not extend their proboscis like honeybees (Daly and Smith, 2000). EMG signal was filtered 0.1 Hz to 1 KHz and amplified 100 times using a differential amplifier (Warner Instruments, Hamden, CT, USA). We scored each trial using amplified EMG signal played through a loud speaker. Pilot data suggested that our moths required ca. 10 trials to associate odor arrival side with a sucrose reward. Initially, moths were given 10-12 trials of conditioning stimuli i.e., odor was presented for 5 s to one of the antennae, and 3 s after odor stimulus onset, they were rewarded with 3 µL of 10 % sucrose solution (*see Supplementary document*). After training, moths were tested with trials in which odor arrived either on the associated or the unassociated side according to a pseudo-random list. Moths were expected to generate feeding muscle EMGs when odor was presented to the antenna on the associated side, and no muscle activity during odor presentation to the opposite antenna (Fig. 1b). Each trial was followed by an inter-trial interval ranging from 30-60 s randomly decided by the control software. During the test phase, moths received no reward.

### (d) Data analysis

#### Response quantification

Response of the moth was scored based on presence (attempted feeding) or absence (no response) of EMG activity. We sorted the test trials based on odor arrival side and for each odor arrival side, we computed moth’s performance using the following formula:

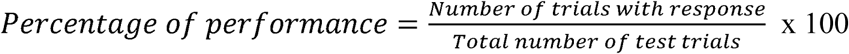

#### Generalized linear mixed-effects model

We analyzed the effect of odor arrival side on the observed behavioral response, i.e., the presence or absence of EMG response by individual moths during the trials and estimated the response variability across individuals using generalized linear mixed-effects model (GLMM) with binomial distribution. We used lme4 package in the statistical software R to analyze the data (Bates et al., 2015; R Core Team, 2019). The data preparation for generalized mixed effect model is explained in the Supplementary document. The GLMM used is as follows:

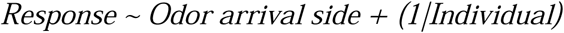

In the model, odor arrival side specified the fixed effect and random effects was specified by (*1|Individual*).

## 3. Results and discussion

Studies on sensory control of behavior routinely deliver experimentally-controlled stimuli and measure the organism’s response (i.e., cell, circuit, or behavior) relative to stimulus onset, offset and/or duration. Most animals studied are bilaterally symmetric, with paired symmetrical sensory structures, but the impacts of their asymmetrical activation is often ignored. In this study, we tested the ability of *M. sexta* to sense odor-arrival direction by stimulating asymmetrically. We show that moths can discriminate odor arrival on one antenna relative to the other using an associative learning test.

The hawkmoth *M. sexta* learned to associate odorant-arrival side with sucrose reward. In our experimental population, the number of moths that learned the linalool and benzaldehyde arrival side with reward were 67.8% (19/28) and 85% (17/20), respectively (Fig.2a&d). We observed that more number of experimental moths learned to associate benzaldehyde with a sugar reward compared to the odorant linalool. Linalool is a common plant odor that is known to induce oviposition behavior in mated female moths and is also a component of floral scent (Bisch-Knaden et al., 2018; Raguso, 2016; Reisenman et al., 2010; Saveer et al., 2012). On the other hand, benzaldehyde could be categorized as a floral scent, as it is emitted by many species of night blooming flowers and is known to act as a pollinator attractant (Raguso et al., 2003a; Raguso et al., 2003b). It may be that *M. sexta* females are more likely to associate the ecologically relevant conditioning stimulus of a sucrose meal with a known floral scent like benzaldehyde, than the known oviposition stimulus linalool. Further studies are required to resolve that question.

Moths showed high performance, >70% EMG response (left: 74.2 ± 5.6%; right: 77.7 ± 9.65%) when they received odor associated linalool arriving on a specific side with the reward when tested. The same moths performed poorly (left: 21.3 ± 8%; right: 23.7 ± 8.9%) when linalool was delivered to the unassociated side (Fig. 2b; Left n = 9; Fig. 2c; Right n = 10). GLMM analysis (*see Materials and Methods & Supplementary document*) showed a strong effect of odor arrival side on moth’s response, with negligible variability between individuals (Table 1). The estimated EMG response probabilities were higher (0.75 & 0.74) when linalool was presented on the same side as the associated side, and lower (0.27 & 0.28) when presented on the unassociated side (see *Supplementary document*).

**Table 1.** Summary of generalized linear mixed effect model for linalool arrival side discrimination data set.

Model: *Response* ~ *Linalool arrival side* + (*1/Individual*)
|  | **Fixed effects** | **Log-odds of response (95% CI)** | **z value** | **Random effects** | **Variance ± (SD)** |
| --- | --- | --- | --- | --- | --- |
| Number of individuals = 9<br>Number of trials = 301 | Associated side (Left) | 1.026 (0.655 to 1.396) | 5.426*** | Individual | 0 |
|  | Unassociated side (Right) | -1.969 (-2.477 to -1.460) | -7.589*** |  |  |
| Number of individuals = 10<br>Number of trials = 369 | Associated side (Right) | 1.119 (0.796 to 1.441) | 6.799*** | Individual | 0 |
|  | Unassociated side (Left) | -2.111 (-2.578 to -1.643) | -8.849*** |  |  |

\*\*\* $P < 0.0001$

**Figure 2.**
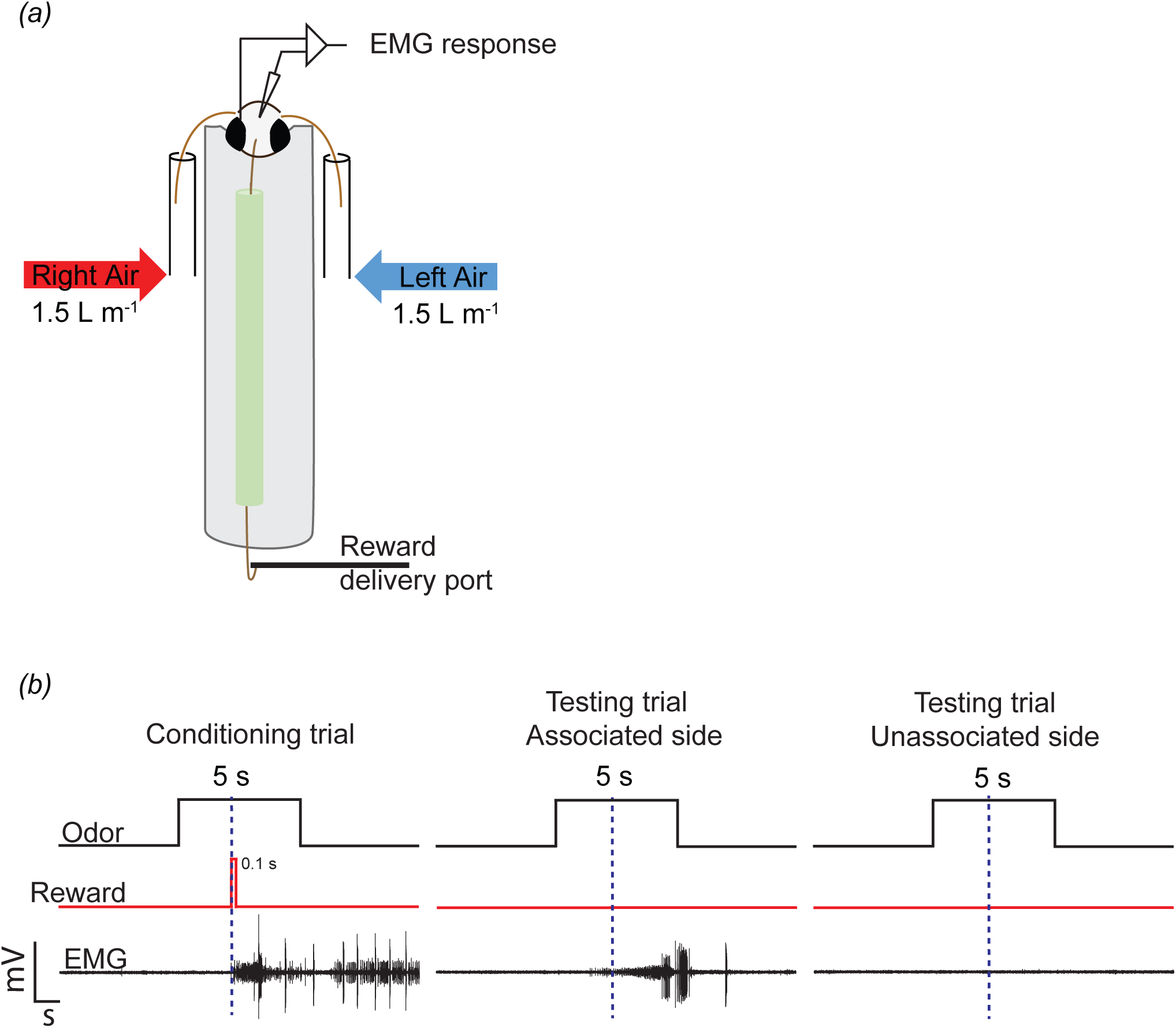
Odor-arrival side discrimination response. Pie charts show the number of moths that learned the odor-arrival side task for both linalool (a) and benzaldehyde (d). Bar graphs showing EMG % response for both left (blue bar) and right (red bar) side odor delivery. Panel b & e show moths response when left antenna was conditioned, and panels c & f when right antenna was conditioned. (b & c) Moths discriminated linalool arrival side. Response to the associated side was higher, > 70% compared to the unassociated side, < 30%. (c) Moths discriminated benzaldehyde arrival side. Response to the associated side was higher, > 70% compared to the unassociated side, < 25%. (b, c, e & f) Error bars indicate standard error of mean.

We observed a similar trend, when we tested moths that associated benzaldehyde arrival on a specific antenna with reward. When moths experienced benzaldehyde on the associated side, they showed EMG response for more than 70% of stimulus presentations (left: 74.8±5.4%; right: 78.4±7.2%). While the same odor was given to unassociated side, the moths response was poor as they responded only to 16 ± 6.6% of left and 14.7 ± 6.8% right side stimulus presentations (Fig. 2e, left n = 8; Fig. 2f, Right n = 9). Using GLMM analysis, we again determined the effects of benzaldehyde arrival side on moth’s response. The odorant arrival side strongly contributed to moth’s response (Table 2) and the estimated probabilities (*see supplementary document*) were high (0.70 & 0.76) for associated side and low for unassociated side (0.16 & 0.15). GLMM showed that performance variability between individuals in odor-arrival side discrimination task was negligible (Table 2).

**Table 2.** Summary of generalized linear mixed effect model for benzaldehyde arrival side discrimination data.

| Model: *Response ~ Benzaldehyde arrival side + (1/Individual)* |  |  |  |  |  |
| --- | --- | --- | --- | --- | --- |
| | Fixed effects | Log-odds of response (95% CI) | z value | Random effects | Variance ± (SD) |
| Number of individuals = 8<br>Number of trials = 185 | Associated side (Left) | 1.099 (0.626 to 1.570) | 4.563*** | Individual | 1.775e-16 (1.332e-08) |
|  | Unassociated side (Right) | -2.747 (-3.474 to -2.021) | -7.410*** |  |  |
| Number of individuals = 9<br>Number of trials = 236 | Associated side (Right) | 1.176 (0.759 to 1.592) | 5.536*** | Individual | 0 |
|  | Unassociated side (Left) | -2.907 (-3.569 to -2.244) | -8.597*** |  |  |

\*\*\* $P < 0.0001$

Animals with bilaterally symmetric odor sensors, i.e., antennae in insects and nostrils in mammals, have been shown to take more time to locate the source and their probability to locate the odor sources reduces, compared to animals with only one sensor (Vickers and Baker, 1991; Sanders and Lucuik, 1992; Lockey and Willis, 2015; Rajan et al., 2006; Porter et al., 2007; Khan et al., 2012; Catania, 2013; Kraus-Epley and Moore, 2002). The loss of one of the sensors may not only reduce the received odor input it is also likely to remove spatial representation of odor information. How moths could use spatial odor information during odor-guided behaviors, such as foraging or mate finding are discussed in the following section.

The ability to extract odor direction information during plume tracking could prove to be useful especially when the cross-section of the odor plume becomes narrow near the source. In addition, the moths slow down and transition to hovering flight as they approach the odor source. Near the source, even small lateral movements of the moths’ flight trajectory may introduce a bias in the amount of odor experienced by each antennae. Since the antennal tips are held ca. 2 cm apart in flight, one of the antennae is likely to be either fully or partially outside the plume. This asymmetrical odor experience would not only inform the moth about loss of odor in one the antennae but also about its position relative to the plume boundary. When foraging in a floral patch, moth’s choice of flower to feed nectar from may depend on the spatial odor information i.e., right or left, along with visual cues. This directional information could be extracted by comparing the odor inputs received across the antennae.

Such a bilateral comparison would require the laterality of the bilateral odor information to be maintained until it reaches the higher brain region where bilateral information is processed. Neural recordings from the lateral accessary lobe (LAL) region of the hawkmoth (*Manduca sexta*) brain showed neurons responding to ipsilateral odor stimulation, suggesting that laterality of the odor information is maintained in higher brain regions (Kanzaki et al., 1991). Additionally, both left and right LALs are connected via commissure of lateral accessory lobe and bilateral odor information is thought to be integrated in the LALs (Homberg et al., 1988; Kanzaki et al., 1991).

Taken together, *M. sexta* moths likely increase their use of bilaterally sampled odor distribution in the near-source plume to locate the odor source. We demonstrate here that *M. sexta* can discriminate between asymmetric odorant onsets between its antennae, thereby opening several questions about odor modulation of plume tracking that may have previously escaped our notice.

## Ethics

### Data accessibility

We have uploaded datasets (CSV files) and R script for generalized linear mixed model analysis.

## Authors’ contribution

K.P. designed and performed research. K.P and M.A.W analyzed data and wrote the paper. Funding acquired by M.A.W.

## Competing interests

### Funding

This work was supported by the Air Force Office of Scientific Research (FA9550-12-1-0237 and

FA9550-14-1-0398 to M.A.W.).

## Acknowledgements

We thank Dr. Kevin Daly for suggestions on PER conditioning setup. Alison Smith and Catherine Lange for helping with data collection and Kim Thompson for rearing experimental animals. We thank Dr. Sanjay Sane for his helpful comments on the manuscript.

## References

Arbas, E. A., Willis, M. A. and Kanzaki, R. (1993). Organization of goal-oriented locomotion: pheromone-modulated flight behavior of moths. In Biological neural networks in invertebrate neuroethology and robotics. (ed. Beer, R. D.), Ritzmann, R. E.), and McKenna, T. M.), p. Academic Press.

Baker, C. F. and Montgomery, J. C. (1999). The sensory basis of rheotaxis in the blind Mexican cave fish, Astyanax fasciatus. J Comp Physiol A 184, 519–527.

Bates, D., Mächler, M., Bolker, B. and Walker, S. (2015). Fitting Linear Mixed-Effects Models Using lme4. Journal of Statistical Software 67, 1–48.

Bisch-Knaden, S., Dahake, A., Sachse, S., Knaden, M. and Hansson, B. S. (2018). Spatial Representation of Feeding and Oviposition Odors in the Brain of a Hawkmoth. Cell Reports 22, 2482–2492.

Borst, A. and Heisenberg, M. (1982). Osmotropotaxis in Drosophila melanogaster. J. Comp. Physiol. 147, 479–484.

Cardé, R. T. and Willis, M. A. (2008). Navigational strategies used by insects to find distant, wind-borne sources of odor. J. Chem. Ecol. 34, 854–866.

Catania, K. C. (2013). Stereo and serial sniffing guide navigation to an odour source in a mammal. Nat Commun 4, 1441.

Daly, K. C. and Smith, B. H. (2000). Associative olfactory learning in the moth Manduca sexta. J Exp Biol 203, 2025–2038.

Daly, K. C., Chandra, S., Durtschi, M. L. and Smith, B. H. (2001). The generalization of an olfactory-based conditioned response reveals unique but overlapping odour representations in the moth Manduca sexta. J Exp Biol 204, 3085–3095.

Daly, K. C., Carrell, L. A. and Mwilaria, E. (2008). Characterizing Psychophysical Measures of Discrimination Thresholds and the Effects of Concentration on Discrimination Learning in the Moth Manduca sexta. Chem. Senses 33, 95–106.

Duistermars, B. J., Chow, D. M. and Frye, M. A. (2009). Flies require bilateral sensory input to track odor gradients in flight. Curr Biol 19, 1301–1307.

Dusenbery, D. B. (1992). Sensory ecology7□: how organisms acquire and respond to information. New York□: W.H. Freeman.

Emanuel, M. E. and Dodson, J. J. (1979). Modification of the Rheotropic Behavior of Male Rainbow Trout (*Salmo gairdneri*) by Ovarian Fluid. J. Fish. Res. Bd. Can. 36, 63–68.

Fraenkel, G. S. and Gunn, D. L. (1961). The orientation of animals: Kineses, taxes and compass reactions. Oxford, England: Dover.

Fry, S. N., Rohrseitz, N., Straw, A. D. and Dickinson, M. H. (2009). Visual control of flight speed in Drosophila melanogaster. Journal of Experimental Biology 212, 1120–1130.

Gardiner, J. M. and Atema, J. (2010). The function of bilateral odor arrival time differences in olfactory orientation of sharks. Curr. Biol. 20, 1187–1191.

Gomez-Marin, A., Stephens, G. J. and Louis, M. (2011). Active sampling and decision making in Drosophila chemotaxis. Nat Commun 2, 441.

Homberg, U., Montague, R. A. and Hildebrand, J. G. (1988). Anatomy of antenno-cerebral pathways in the brain of the sphinx moth Manduca sexta. Cell and Tissue Research 254,.

Kanzaki, R., Arbas, E. A. and Hildebrand, J. G. (1991). Physiology and morphology of protocerebral olfactory neurons in the male moth Manduca sexta. J Comp Physiol A 168, 281–298.

Kennedy, J. S. (1940). The Visual Responses of Flying Mosquitoes. Proceedings of the Zoological Society of London A109, 221–242.

Kennedy, J. S. (1983). Zigzagging and casting as a programmed response to wind-borne odour: a review. Physiological Entomology 8, 109–120.

Kennedy, J. S. and Marsh, D. (1974). Pheromone-Regulated Anemotaxis in Flying Moths. Science 184, 999–1001.

Khan, A. G., Sarangi, M. and Bhalla, U. S. (2012). Rats track odour trails accurately using a multi-layered strategy with near-optimal sampling. Nature Communications 3, 703.

Kraus-Epley, K. E. and Moore, P. A. (2002). Bilateral and Unilateral Antennal Lesions Alter Orientation Abilities of the Crayfish, Orconectes rusticus. Chem Senses 27, 49–55.

Kulpa, M., Bak-Coleman, J. and Coombs, S. (2015). The lateral line is necessary for blind cavefish rheotaxis in non-uniform flow. Journal of Experimental Biology 218, 1603–1612.

Kuwabara, M. (1957). Bildung des bedingten reflexes von Pavlovs typus bei der honigbiene, Apis mellifica (mit 1 textabbildung). Journal of the Faculty of Science Hokkaido University, Zoology 13, 458–464.

Lockey, J. K. and Willis, M. A. (2015). One antenna, two antennae, big antennae, small: total antennae length, not bilateral symmetry, predicts odor-tracking performance in the American cockroach Periplaneta americana. Journal of Experimental Biology 218, 2156–2165.

Louis, M., Huber, T., Benton, R., Sakmar, T. P. and Vosshall, L. B. (2008). Bilateral olfactory sensory input enhances chemotaxis behavior. Nature Neuroscience 11, 187–199.

Martin, H. (1965). Osmotropotaxis in the Honey-Bee. Nature 208, 59.

Martin, J. P., Beyerlein, A., Dacks, A. M., Reisenman, C. E., Riffell, J. A., Lei, H. and Hildebrand, J. G. (2011). The neurobiology of insect olfaction: sensory processing in a comparative context. Prog. Neurobiol. 95, 427–447.

Page, J. L., Dickman, B. D., Webster, D. R. and Weissburg, M. J. (2011). Staying the course: chemical signal spatial properties and concentration mediate cross-stream motion in turbulent plumes. Journal of Experimental Biology 214, 1513–1522.

Parthasarathy, K. and Bhalla, U. S. (2013). Laterality and Symmetry in Rat Olfactory Behavior and in Physiology of Olfactory Input. Journal of Neuroscience 33, 5750–5760.

Porter, J., Craven, B., Khan, R. M., Chang, S.-J., Kang, I., Judkewitz, B., Volpe, J., Settles, G. and Sobel, N. (2007). Mechanisms of scent-tracking in humans. Nature Neuroscience 10, 27–29.

R Core Team (2019). R: A Language and Environment for Statistical Computing. Vienna, Austria: R Foundation for Statistical Computing.

Raguso, R. A. (2016). More lessons from linalool: insights gained from a ubiquitous floral volatile. Current Opinion in Plant Biology 32, 31–36.

Raguso, R. A., Levin, R. A., Foose, S. E., Holmberg, M. W. and McDade, L. A. (2003a). Fragrance chemistry, nocturnal rhythms and pollination “syndromes” in Nicotiana. Phytochemistry 63, 265–284.

Raguso, R. A., Henzel, C., Buchmann, S. L. and Nabhan, G. P. (2003b). Trumpet Flowers of the Sonoran Desert: Floral Biology of Peniocereus Cacti and Sacred Datura. International Journal of Plant Sciences 164, 877–892.

Rajan, R., Clement, J. P. and Bhalla, U. S. (2006). Rats Smell in Stereo. Science 311, 666–670.

Reisenman, C. E., Riffell, J. A., Bernays, E. A. and Hildebrand, J. G. (2010). Antagonistic effects of floral scent in an insect-plant interaction. Proc. Biol. Sci. 277, 2371–2379.

Riffell, J. A., Lei, H., Christensen, T. A. and Hildebrand, J. G. (2009). Characterization and Coding of Behaviorally Significant Odor Mixtures. Current Biology 19, 335–340.

Sanders, C. J. and Lucuik, G. S. M. (1992). MATING BEHAVIOR OF SPRUCE BUDWORM MOTHS, CHORISTONEURA FUMIFERANA (CLEM.) (LEPIDOPTERA: TORTRICIDAE). The Canadian Entomologist 124, 273–286.

Sasaki, M. and Riddiford, L. M. (1984). Regulation of reproductive behaviour and egg maturation in the tobacco hawk moth, Manduca sexta. Physiological Entomology 9, 315–327.

Saveer, A. M., Kromann, S. H., Birgersson, G., Bengtsson, M., Lindblom, T., Balkenius, A., Hansson, B. S., Witzgall, P., Becher, P. G. and Ignell, R. (2012). Floral to green: mating switches moth olfactory coding and preference. Proc Biol Sci 279, 2314–2322.

Saxena, N., Natesan, D. and Sane, S. P. (2017). Odor source localization in complex visual environments by fruit flies. Journal of Experimental Biology jeb.172023.

Schone, H. and Strausfeld, C. (1984). Spatial Orientation: The Spatial Control of Behavior in Animals and Man. Princeton, New Jersey: Princeton University Press.

Takasaki, T., Namiki, S. and Kanzaki, R. (2012). Use of bilateral information to determine the walking direction during orientation to a pheromone source in the silkmoth Bombyx mori. J. Comp. Physiol. A Neuroethol. Sens. Neural. Behav. Physiol. 198, 295–307.

Vickers, N. J. and Baker, T. C. (1991). The effects of unilateral antennectomy on the flight behaviour of male Heliothis virescens in a pheromone plume. Physiological Entomology 16, 497–506.

Vonbekesy, G. (1964). OLFACTORY ANALOGUE TO DIRECTIONAL HEARING. J Appl Physiol 19, 369–373.

Wasserman, S., Lu, P., Aptekar, J. W. and Frye, M. A. (2012). Flies dynamically anti-track, rather than ballistically escape, aversive odor during flight. Journal of Experimental Biology 215, 2833–2840.

Weissburg, M. J. and Dusenbery, D. B. (2002). Behavioral observations and computer simulations of blue crab movement to a chemical source in a controlled turbulent flow. Journal of Experimental Biology 205, 3387–3398.

Willis, M. A. and Arbas, E. A. (1991). Odor-modulated upwind flight of the sphinx moth, Manduca sexta L. J. Comp. Physiol. A 169, 427–440.

